# Tacrolimus rescues endothelial ALK1 loss-of-function signaling and improves HHT vascular pathology

**DOI:** 10.1101/137737

**Authors:** Santiago Ruiz, Pallavi Chandakkar, Haitian Zhao, Julien Papoin, Prodyot K. Chatterjee, Erica Christen, Christine N. Metz, Lionel Blanc, Fabien Campagne, Philippe Marambaud

## Abstract

Hereditary hemorrhagic telangiectasia (HHT) is a genetic vascular disorder arising from endothelial cell (EC) proliferation and hypervascularization, for which no cure exists. Because HHT is caused by loss-of-function mutations in BMP9-ALK1-Smad1/5/8 signaling, interventions aimed at activating this pathway are of therapeutic value. By screening FDA-approved drug libraries, we identified tacrolimus (FK-506) as a potent activator of Smad1/5/8 in BMP9-challenged reporter cells. In primary ECs, tacrolimus activated Smad1/5/8 to oppose the pro-angiogenic gene expression signature associated with ALK1 loss-of-function, by notably reducing Dll4 expression. In these cells, tacrolimus also inhibited Akt and p38 stimulation by VEGF. In the BMP9/10-immunodepleted postnatal retina—a mouse model of HHT vascular pathology—tacrolimus activated endothelial Smad1/5/8 and prevented the Dll4 overexpression and hypervascularization associated with this model. Finally, tacrolimus stimulated Smad1/5/8 in cells transfected with BMP9-unresponsive ALK1 HHT mutants and in HHT patient blood outgrowth ECs. We propose that tacrolimus repurposing has therapeutic potential in HHT.

## Introduction

Hereditary hemorrhagic telangiectasia (HHT, or Rendu-Osler-Weber syndrome) is an autosomal dominant genetic disease affecting ∼1 in 5,000 individuals. The clinical presentation of HHT includes potentially hemorrhagic vascular anomalies in multiple tissues and organs in the form of arteriovenous malformations (AVMs) and mucocutaneous telangiectasias. The systemic manifestations of HHT make patient management challenging and can lead to highly debilitating and life-threatening hemorrhagic events and secondary cerebral, hepatic, pulmonary, and cardiac complications (1, 2).

Mutations in the genes *ENG* (encoding endoglin) or *ACVRL1* (activin receptor-like kinase 1, ALK1) are the main cause of HHT, and define the two disease subtypes: HHT1 and HHT2, respectively (3, 4). Mutations in *SMAD4* (encoding Smad4) and *GDF2* (bone morphogenetic protein 9, BMP9) cause rare forms of the disease called juvenile polyposis/HHT combined syndrome and HHT-like vascular anomaly syndrome, respectively (3). BMP9, ALK1, endoglin, and Smad4 functionally interact in the same signaling pathway. ALK1 is a BMP type I receptor of the transforming growth factor-β (TGF-β) superfamily, which forms complexes with a BMP type II receptor (e.g., BMPR2) and the co-receptor endoglin. Of note, mutations in BMPR2 cause familial pulmonary arterial hypertension (PAH), a separate clinical entity that is observed in some HHT2 patients (5). ALK1 receptors are abundantly expressed by endothelial cells (ECs) and are specifically activated by the circulating ligands BMP9 and BMP10 (6-9). Once activated, ALK1 receptors phosphorylate the signal transducers Smad1, 5, and 8 to trigger the formation of Smad1/5/8-Smad4 complexes. Smad1/5/8-Smad4 complexes then translocate to the nucleus to control specific gene expression programs (10-12).

HHT mutations cause a loss-of-function in the ALK1 signaling pathway. Indeed, HHT-causing mutations in ALK1 or endoglin block Smad1/5/8 signaling by BMP9 (13-15). Studies in mouse and zebrafish models further revealed that ALK1 signaling inactivation leads to robust vascular defects that include vascular hyperproliferation and AVMs (16-20). The exact cellular processes leading to AVM development in HHT, i.e., the formation of direct shunts between arteries and veins, remain poorly understood. Solid evidence suggests, however, that HHT is caused by abnormal reactivation of angiogenesis (21, 22) and that inhibition of the pro-angiogenic guidance cue, vascular endothelial growth factor (VEGF), might reduce the pathology in HHT mouse models and HHT patients (21, 22), but see also (23, 24).

During angiogenesis, ECs engage in specific gene expression programs enabling them to migrate and proliferate to expand a vascular sprout and ultimately form a new vascular bed structured around arteries connecting with veins *via* the capillaries (25). Angiogenic sprouting is initiated by the activation of VEGF receptor 2 (VEGFR2) by VEGF in a specific subset of ECs. Communication between ECs during angiogenesis is primarily controlled by Dll4/Notch signaling, where VEGFR2 activation triggers Dll4 expression (26). ALK1 receptors interact with Notch signaling during angiogenesis by activating Notch transcriptional targets and by supporting sprouting angiogenesis and EC specification (17-19, 27, 28), but see also (29). Thus, ALK1 is a key regulator of VEGF/Dll4/Notch signaling and EC functions during angiogenesis and vascular maintenance. It is therefore not surprising that ALK1 loss-of-function in HHT is sufficient to cause significant defects in EC integrity that ultimately lead to vascular pathology.

In this study, by using RNA-Seq analyses in human umbilical vein endothelial cells (HUVECs), we show that ALK1 signaling inhibition was associated with a specific transcriptional signature that significantly increased the gene expression of several key pro-angiogenic markers, such as Dll4. By screening ∼700 FDA-approved drugs, we identified tacrolimus as a potent activator of endothelial Smad1/5/8 signaling. We found that tacrolimus significantly reversed the transcriptional signature associated with ALK1 loss-of-function, by notably opposing the gene expression upregulation of the identified pro-angiogenic markers, including Dll4. In addition, tacrolimus reduced the vascular pathology of an HHT mouse model and potently activated Smad1/5/8 signaling in HHT patient-derived primary ECs. These data show that tacrolimus has the potential to block HHT pathology and restore EC homeostasis by rescuing the signaling and gene expression control defects caused by ALK1 loss-of-function.

## Results

### ALK1 inhibition controls a pro-angiogenic gene expression response

To gain insights into the global transcriptional deregulations caused by ALK1 signaling inhibition, we profiled gene expression by RNA-Seq in HUVECs treated with ALK1 extracellular domain-derived ligand trap (ALK1-Fc). Incubation with ALK1-Fc for 24h led to an inhibition of ALK1 signaling, which manifested by a decrease of Smad1/5/8 phosphorylation (pSmad1/5/8) and ID1 (inhibitor of differentiation 1) expression (Fig. 1A, inset). ALK1-Fc treatment significantly changed the expression of 251 genes (*n* = 6 biological replicates per group, Limma Voom test controlling for sample batches, FDR ≤ 0.01; Figs. 1A and 1B, and Table S1). Among the up-regulated genes, several had previously been described as pro-angiogenic and/or endothelial tip cell markers (26, 30-33), including *DLL4*, *ANGPT2*, *KDR*, *CXCR4*, *PGF*, and *SMOC1* (Figs. 1A and 1B, and Table S1). Dll4 is of particular interest because of its central role in angiogenesis (26). We confirmed at the protein level that ALK1 inhibition by ALK1-Fc robustly and significantly increased Dll4 expression by HUVECs (Figs. 2A and 2B). Importantly, treatment with the physiological activating ALK1 ligand, BMP9, led to the opposite effect, by significantly reducing Dll4 expression (Figs. 2A and 2B). These whole-transcriptome and protein expression analyses are consistent with the concept that ALK1 inhibition is associated with a pro-angiogenic gene expression program.

**Figure 1.**
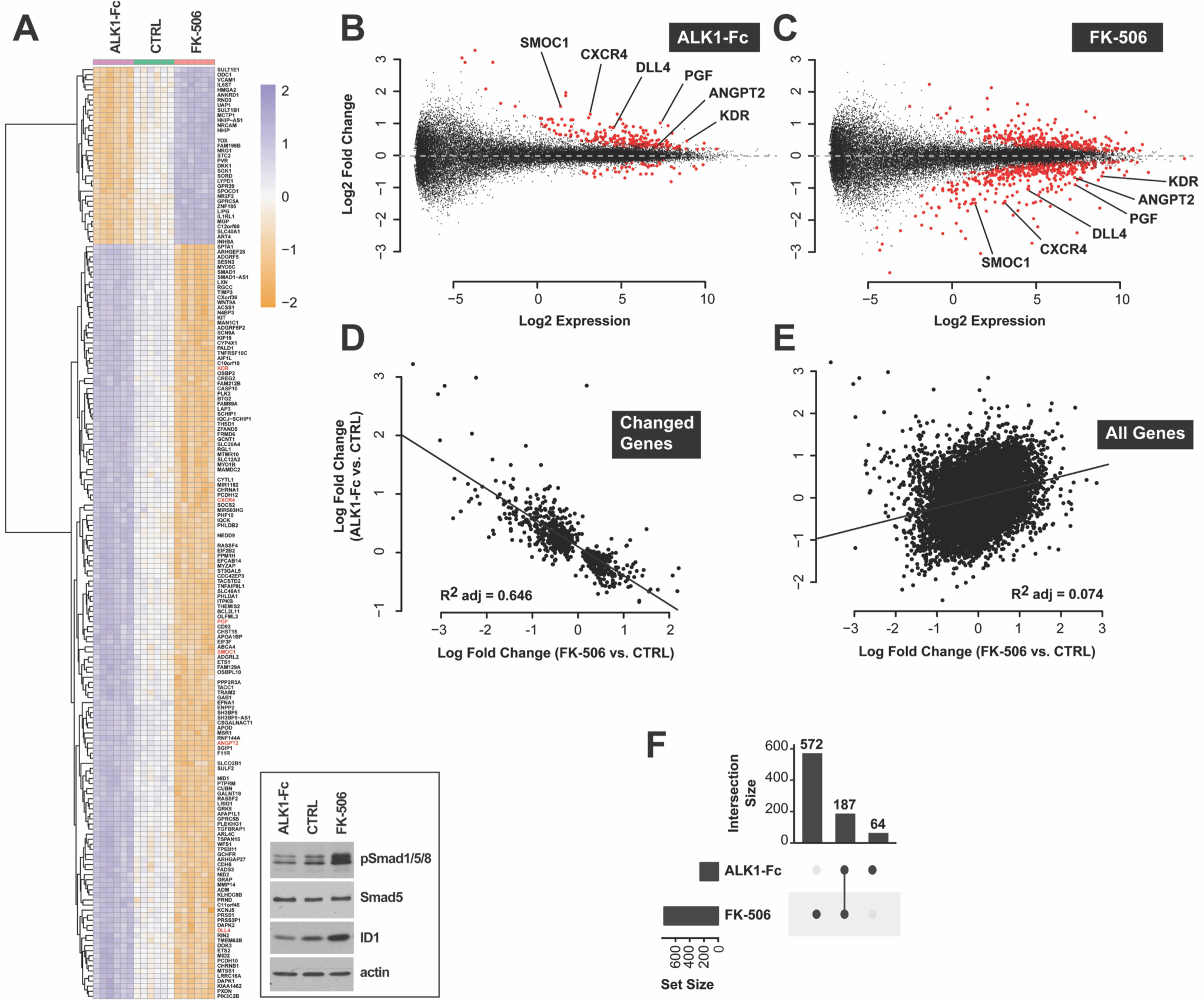
Tacrolimus mimics ALK1-mediated gene expression control. (**A**) RNA-Seq heat map of gene expression changes in HUVECs treated or not (CTRL) for 24h in complete medium (conditioned for 2 days) with ALK1-Fc (1 μg/mL) or tacrolimus (FK-506, 0.3 μM) (*n* = 6). Inset, cell extracts were analyzed by Western blotting (WB) using antibodies directed against the indicated proteins. (**B and C**) RNA-Seq Mean Average Plots (MA-Plots) displaying all differentially expressed transcripts in ALK1-Fc-treated (B) and FK506-treated (C) HUVECs. Genes with a FDR < 0.01 are shown in red. Genes unchanged by treatment are expected to lie on the horizontal dashed line at log2 fold change = 0. Low expression values reflect experimental measurement noise. Genes discussed in the text were annotated on the plots. (**D and E**) RNA-Seq scatter plots comparing gene expression changes (log fold change) between ALK1-Fc and FK-506 treatments for transcripts differentially expressed by either treatment (FDR < 0.01, D) and for all transcripts (E). Plot in (D) shows a clear inverse correlation and indicates that FK-506 treatment had the inverse effect of ALK1-Fc. (**F**) RNA-Seq UpSet plot showing the size of set intersections for transcripts differentially expressed by ALK1-Fc or FK506 treatments. Bars indicate how many transcripts were differentially expressed in each labeled condition. Lines connect dots to indicate set intersection. For instance, 572+187 transcripts were differentially expressed by FK-506 treatment, of which 187 were also differentially expressed when HUVECs were treated with ALK1-Fc. 64 genes were differentially expressed following ALK1-Fc treatment, but not in the other condition.

**Figure 2.**
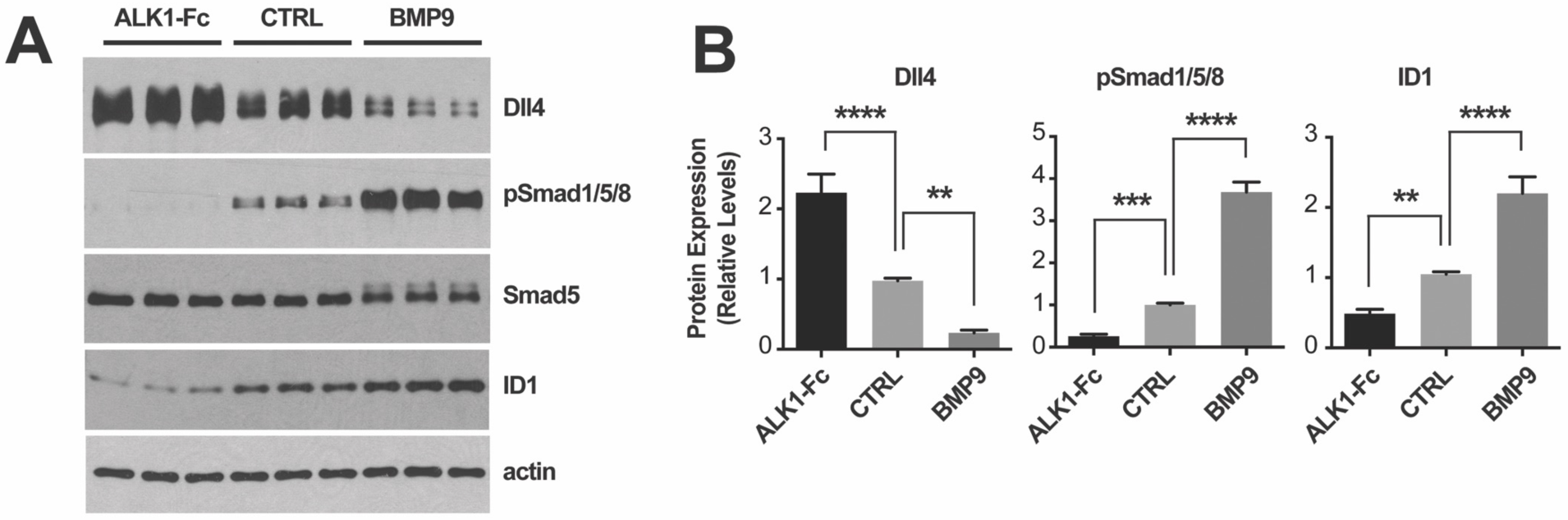
ALK1 signaling controls Dll4 expression. (**A**) HUVECs were treated or not (CTRL) for 24h in complete medium (conditioned for 2 days) with ALK1-Fc (1 μg/mL) or BMP9 (10 ng/mL). Cell extracts were analyzed by WB using antibodies directed against the indicated proteins. (**B**) Densitometric analyses and quantification of Dll4, pSmad1/5/8, and ID1 relative levels in *n* = 4 independent experiments, as in (A). Data represent mean ± s.e.m.; \*\**P* < 0.01, \*\*\**P* < 0.001, \*\*\*\**P* < 0.0001; one-way ANOVA, Dunnett’s multiple comparisons test.

### Tacrolimus is a potent Smad1/5/8 signaling activator

We next screened the NIH clinical collections (NCCs) of ∼700 FDA-approved drugs for their potential to activate Smad1/5/8 signaling in reporter C2C12 myoblast cells expressing luciferase under the control of an *Id1* promoter response element [C2C12BRA cells (34)]. This approach previously identified BMPR2 signaling activators in BMP4-challenged reporter cells (35). Here, we asked whether we could identify drugs that enhance BMP9-specific Smad1/5/8 signaling. To this end, we screened the NCC drugs in C2C12BRA cells maintained in medium depleted of exogenous growth factors and supplemented with a half-maximal effective concentration (EC_50_) of BMP9. We determined that 0.5 ng/mL was the EC_50_ of BMP9 in C2C12BRA cells when the cells were challenged for 24h in depleted medium (0.1% FBS, Fig. S1A). NCC drugs were thus screened using C2C12BRA cells maintained in depleted medium supplemented with 0.5 ng/mL BMP9 for 24h. The most potent activating drug was tacrolimus (FK-506; fold change = 3.4, Z score = 9.7; Fig. 3A). The effect of tacrolimus in C2C12BRA cells was confirmed using another commercial source of the drug, and the EC_50_ of tacrolimus was determined to be of 37 nM in depleted medium supplemented with BMP9 at EC_50_ (Fig. 3B) and of 85 nM in complete culture medium containing 10% FBS (Fig. S1B). At the protein level and in a dose-dependent manner, tacrolimus increased ID1 levels by C2C12 cells (Fig. 3C) and HUVECs (Fig. 3D). Importantly, tacrolimus also efficiently elevated pSmad1/5/8 in both C2C12 cells and HUVECs (Figs. 3C and 3D). Thus, tacrolimus is a potent trigger of Smad1/5/8 signaling in ECs under conditions of specific challenge with BMP9 and in complete medium containing regular serum exogenous trophic factors. These data are in line with the previously reported activating effect of tacrolimus on BMPR2-Smad1/5/8 signaling in pulmonary ECs (35). Thus, tacrolimus is a potent activator of the BMP9-ALK1-Smad1/5/8-ID1 signaling cascade.

**Figure 3.**
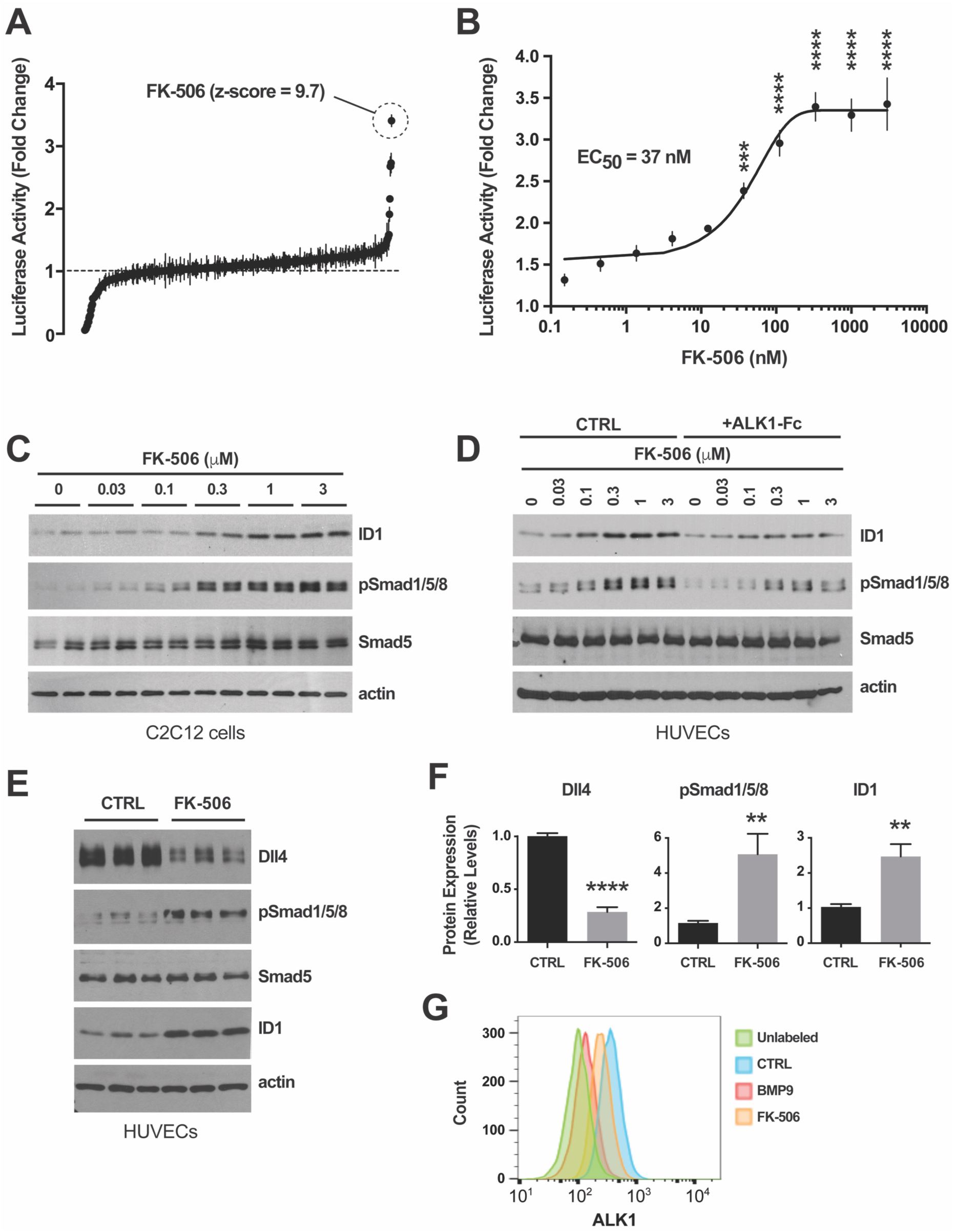
Identification and validation of tacrolimus as a Smad1/5/8 signaling activator. (**A**) NCC screen using *Id1* transactivation luciferase assay in C2C12BRA cells. Drugs were tested in duplicate at 3 μM on cells maintained for 24h in depleted medium containing 0.1% FBS and 0.5 ng/mL BMP9 (BMP9 EC_50_). Data (fold change mean ± range) are ranked by mean score and were normalized to DMSO control. (**B**) Luciferase activity in C2C12BRA cells treated with different concentrations of tacrolimus (FK-506) in depleted medium supplemented with BMP9 at EC_50,_ as in (A). Data represent mean ± s.e.m. (*n* = 4); \*\*\**P* < 0.001, \*\*\*\**P* < 0.0001; one-way ANOVA, Bonferroni’s multiple comparisons test. (**C-E**) C2C12 cells (C) and HUVECs (D and E) were treated or not (CTRL) for 24h in complete medium (conditioned for 2 days) with ALK1-Fc (1 μg/mL, D) or tacrolimus at the indicated concentrations (C and D) or 0.3 μM (E). Cell extracts were analyzed by WB using antibodies directed against the indicated proteins. (**F**) Densitometric analyses and quantification of Dll4, pSmad1/5/8, and ID1 relative levels in *n* = 3 independent experiments, as in (E). Data represent mean ± s.e.m.; \*\**P* < 0.01, \*\*\*\**P* < 0.0001; Student’s *t*-test. (**G**) Flow cytometric analysis of surface ALK1 expression in HUVECs treated or not (CTRL) for 24h in complete medium (conditioned for 2 days) with BMP9 (10 ng/mL) or tacrolimus (0.3 μM).

### Tacrolimus rescues the gene expression deregulations associated with ALK1 inhibition

At the whole-transcriptome level, we compared the effect of tacrolimus and ALK1-Fc on gene expression in HUVECs. RNA-Seq revealed that Smad1/5/8 signaling activation by tacrolimus (Fig. 1A, inset) significantly altered the expression of 759 genes (FDR ≤ 0.01, Figs. 1A and 1C, and Table S1). Strikingly, 187 of these deregulated genes were also found to be deregulated by ALK1-Fc treatment (Fig. 1F), but in the opposite direction (Figs. 1A-C). Indeed, a significant inverse correlation in fold change (log2) was found for genes either differentially expressed by ALK1-Fc treatment (vs. control) or tacrolimus treatment (vs. control) (FDR < 0.01 either condition, Fig. 1D). The same comparison between all genes did not show a similar correlation (Fig. 1E), validating the specificity of the observed effect. When restricting this intersection to genes differentially expressed in both conditions (changed by both ALK1-Fc and tacrolimus treatments), 187 genes were identified that included *DLL4*, *ANGPT2*, *KDR*, *CXCR4*, *PGF,* and *SMOC1* (Figs. 1A-C, F). In agreement with the effect of BMP9 treatment on Dll4 expression (Fig. 2), Smad1/5/8 signaling activation by tacrolimus robustly and significantly reduced Dll4 protein expression by HUVECs (Figs. 3E and 3F).

To provide further evidence that tacrolimus activates Smad1/5/8 signaling and to increase our understanding of the mechanism by which this signaling is activated, we asked whether tacrolimus affects ALK1 trafficking, as TGF-β receptors are internalized upon ligand binding during signal transduction (36). Flow cytometry analyses using an antibody directed against the extracellular region of ALK1 revealed an almost complete loss of the endogenous levels of ALK1 at the cell surface upon BMP9 treatment in HUVECs (Fig. 3G), suggesting that ALK1, like TGF-β receptors, might internalize upon receptor activation. Interestingly, tacrolimus, at a concentration activating Smad1/5/8, also caused a decrease of surface ALK1 (Fig. 3G), indicating that tacrolimus mimics several important aspects of BMP9 activation and that it acts upstream in the signaling cascade at the level of the receptor.

### Tacrolimus inhibits VEGF-mediated AKT and p38 activation

Increased Dll4 expression is a critical transcriptional response to VEGF/VEGFR2 signaling in ECs during angiogenesis (37). We sought to determine whether the repressing effect of ALK1 activation on Dll4 expression could be due to inhibition of VEGFR2 signal transduction. At least three major kinase pathways are activated in ECs upon VEGFR2 signaling: Akt/PKB, p38 MAPK, and Erk1/2. Compelling evidence has highlighted the key role of the PI3K/Akt axis in Dll4 transcriptional activation by VEGF (38, 39). We found that pretreatment with BMP9 for 24h had no effect on Akt, p38, or Erk1/2 phosphorylation upon VEGF stimulation in HUVECs, whereas BMP9 efficiently activated ALK1 signaling by increasing pSmad1/5/8 and ID1 levels under these conditions (Figs. 4A and 4B). Surprisingly, tacrolimus treatment of HUVECs for 24h prior to VEGF stimulation resulted in a complete inhibition of Akt phosphorylation (both at Thr-308 and Ser-473), and in a partial but significant reduction in p38 phosphorylation (Figs. 4A and 4B). The effect of tacrolimus was specific towards Akt and p38 because the drug did not affect VEGF-mediated Erk1/2 phosphorylation (Figs. 4A and 4B). Together, these data show that ALK1 signaling can control Dll4 expression independent of VEGF signaling. Therefore, the potent repressing effect of tacrolimus on Dll4 expression is likely to be due to a dual and independent control of the Smad1/5/8 and VEGF pathways (see model in Fig. 4C).

**Figure 4.**
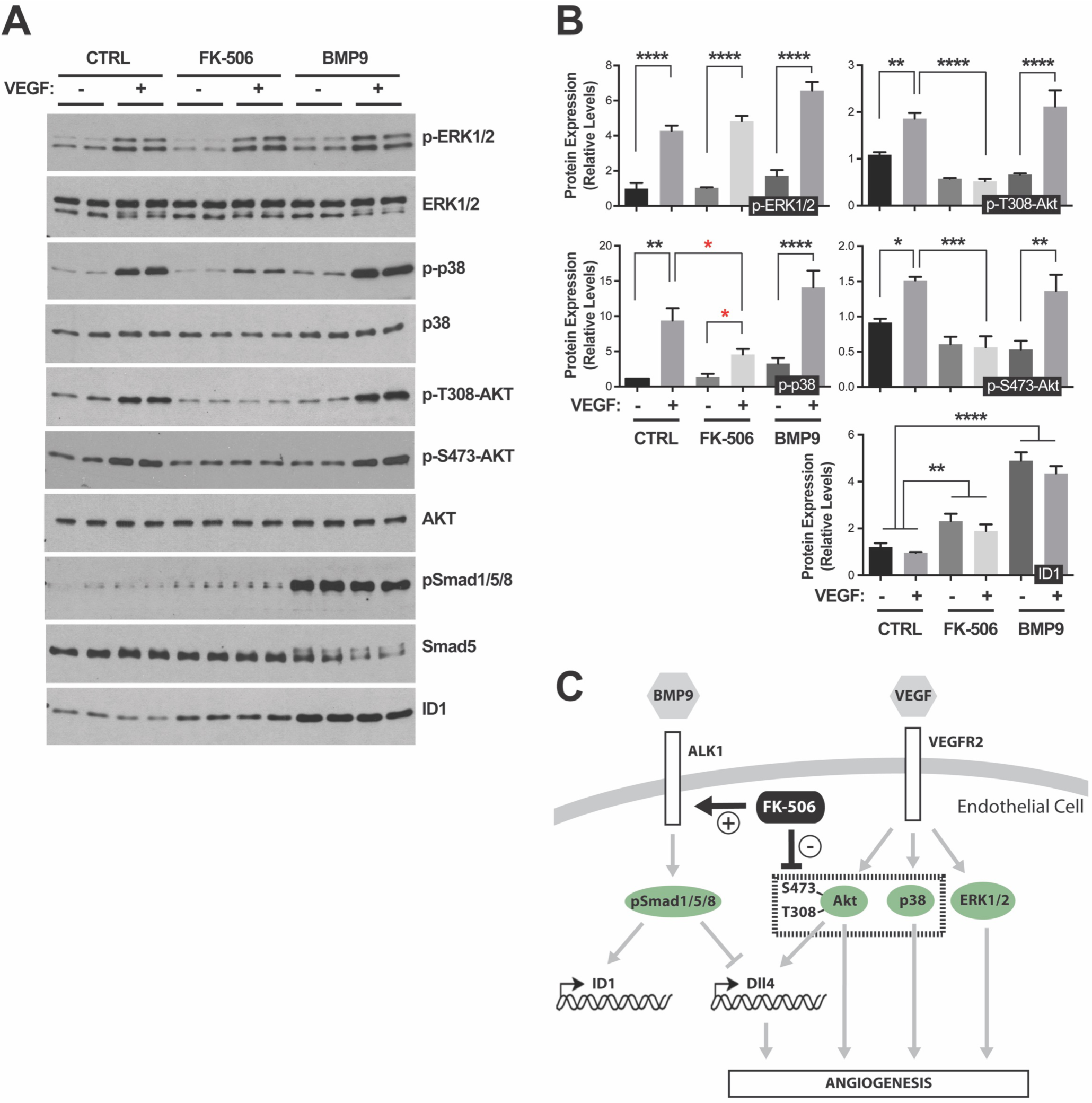
Tacrolimus inhibits VEGF-mediated Akt and p38 activation. (**A**) HUVECs treated or not (CTRL) for 24h (in 0.05% FBS medium) with BMP9 (10 ng/mL) or tacrolimus (FK-506, 0.3 μM), were stimulated for 5 min with VEGF (25 ng/mL). Cell extracts were analyzed by WB using antibodies directed against the indicated proteins. (**B**) Densitometric analyses and quantification of the relative levels of the indicated proteins in *n* = 3 independent experiments, as in (A). Data represent mean ± s.e.m.; \**P* < 0.05, \*\**P* < 0.01, \*\*\**P* < 0.001, \*\*\*\**P* < 0.0001; one-way ANOVA, Dunnett’s and Sidak’s multiple comparisons tests. (**C**) Schematic illustration of the proposed effect of tacrolimus on ALK1 and VEGFR2 signaling pathways.

### Tacrolimus improves vascular pathology in the BMP9/10-immunoblocked retina

Recently, we described a new HHT mouse model, which has the advantage of being practical and robust, as well as particularly suitable and reliable for the analysis of large cohorts (40). In this model, we showed that transmammary delivery of BMP9 and BMP10 blocking antibodies to nursing mouse pups led to the inhibition of ALK1 signaling and development of an HHT-like pathology in their postnatal retinal vasculature. This pathology included the presence of robust hypervascularization and AVM development (40). Using this model, we assessed the *in vivo* potential of tacrolimus to modulate Smad1/5/8 signaling and HHT vascular pathology. Pathology in the newborn retina was triggered by one intraperitoneal (i.p.) injection of anti-BMP9/10 antibodies in lactating dams on postnatal day 3 (P3), as reported before (40). In parallel, pups were treated with tacrolimus (0.5 mg/kg/d, i.p.) from P3 until they were euthanized on P6 to analyze their retinal vasculature. Tacrolimus slightly but significantly decreased both the number of AVMs and their diameter (Figs. 5A and 5B). More strikingly, tacrolimus efficiently reduced the density of the vascular plexus by preventing the appearance of the hyperbranched phenotype caused by BMP9/10 immunoblocking (Figs. 5C-H). We quantified the surface area occupied by the retinal vasculature at the capillary plexus and found that tacrolimus treatment significantly reduced the abnormal vascular density caused by BMP9/10 immunoblocking (Figs. 5F-I).

**Figure 5.**
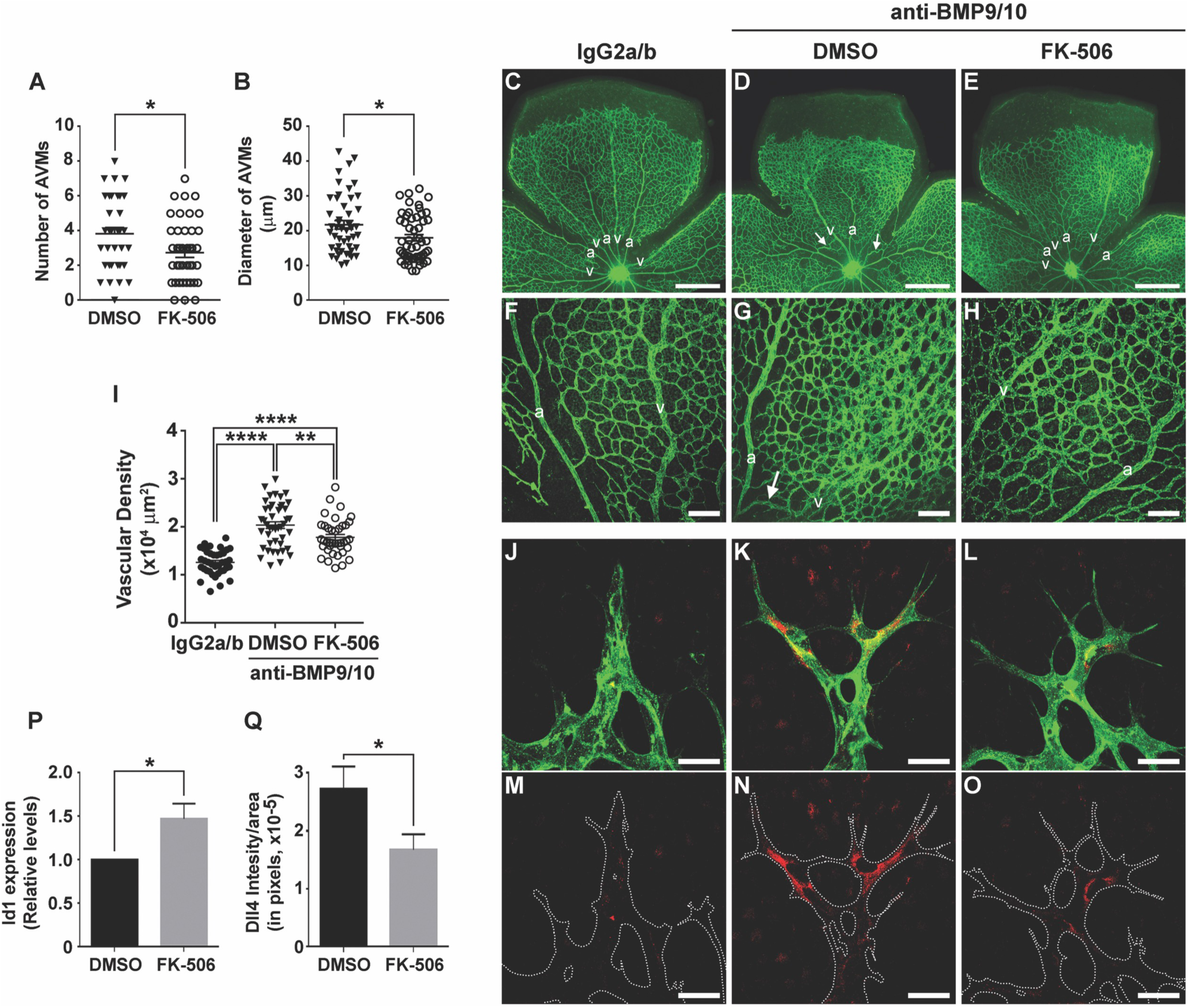
Tacrolimus improves vascular pathology in the BMP9/10-immunoblocked retina. (**A and B**) AVM number (A) and AVM diameter (B) in the retinal vasculature of P6 pups treated *via* the transmammary route with BMP9 and BMP10 blocking antibodies [see Methods and Ref. (40)], and treated with tacrolimus (FK-506, 0.5 mg/kg/d) or vehicle (DMSO). Data represent mean ± s.e.m. per retina (*n* = 10-20 pups per group); \**P* < 0.05; Student’s *t*-test (A) and Mann Whitney U test (B). (**C-H**) Representative images of retinas stained with fluorescent isolectin B4 from pups treated or not (DMSO) with tacrolimus (FK-506, 0.5 mg/kg/d), and treated *via* the transmammary route with control IgG2a/b (C and F) or BMP9/10 blocking antibodies (D, E, G, and H). Higher magnifications in (F-H) show retinal vasculature fields (plexus area) between an artery (a) and a vein (v). Arrows in (D) and (G) denote AVMs. (**I**) Scatter plot showing the density of the retinal vascular plexus in pups treated as in (F-H). Data represent mean ± s.e.m. (*n* = 6-8); \*\**P* < 0.01, \*\*\*\**P* < 0.0001; one-way ANOVA, Tukey’s multiple comparisons test. (**J-O**) Histochemistry analysis of the vascular front of P6 retinas treated as in (C-H) and stained with fluorescent isolectin B4 (J-L, green) and anti-Dll4 antibody (J-O, red). (**P**) Retinal ECs isolated with anti-CD31 microbeads from pups treated as in (A) were analyzed for *Id1* mRNA levels by RT-qPCR. The results are expressed as relative levels of the control condition (*n* = 3 determinations). (**Q**) Quantification of Dll4 levels in 3 experiments as in (N and O). Data in (P) and (Q) represent mean ± (*n* = 3-5); \**P* < 0.05; Student’s *t*-test (P) and Mann Whitney U test (Q). Scale bars, 500 μm (C-E), 100 μm (F-H), 30 μm (J-O).

In agreement with the *in vitro* data using ALK1-Fc in HUVECs (Fig. 2), we further found that ALK1 signaling interference *via* BMP9/10 immunoblocking in mouse pups caused a pronounced increase in Dll4 expression *in vivo* in their retinal ECs at the developing vascular front (Figs. 5J, 5K, 5M, and 5N). Importantly, in retinal ECs, tacrolimus treatment not only increased Smad1/5/8 signaling by elevating *Id1* gene expression (Fig. 5P), but also prevented the Dll4 overexpression caused by BMP9/10 immunoblocking (Figs. 5J-O and 5Q). In summary, tacrolimus activated Smad1/5/8 signaling and reduced Dll4 overexpression *in vivo* in ECs, and inhibited HHT pathology in a mouse model of ALK1 loss-of-function.

### Tacrolimus activates Smad1/5/8 in cells expressing ALK1 HHT mutants

The *in vivo* data obtained in the BMP9/10-immunoblocked retina show that tacrolimus activates endothelial Smad1/5/8 signaling even in the presence of strong ALK1 blockade. In cell cultures, we further found that, although ALK1-Fc treatment reduced the effect of tacrolimus on Smad1/5/8 signaling, it still allowed the drug to strongly increase luciferase activity in C2C12BRA cells (Fig. S1B) and pSmad1/5/8 and ID1 levels in HUVECs (Fig. 3D). These results strongly support the concept that tacrolimus is able to bypass a partial, or even complete, ALK1 loss-of-function caused by mutations in HHT patients. To address this possibility, we investigated the effect of tacrolimus on Smad1/5/8 activation in C2C12 cells transfected with two inactive HHT ALK1 mutants, R411P and T372fsX (14, 41, 42). We found that over-expression of R411P-ALK1 (Fig. 6A) or T372fsX-ALK1 (Fig. 6B) resulted in a nearly complete inhibition of Smad1/5/8 activation by BMP9, when compared to cells overexpressing comparable levels of wild type (WT) ALK1, in which BMP9, as expected, triggered a robust increase of pSmad1/5/8 (Figs. 6A and 6B). In contrast, tacrolimus treatment efficiently activated Smad1/5/8 in cells expressing either WT ALK1 or the ALK1 HHT mutants (Figs. 6A and 6B). To extend our observations and demonstrate efficacy using a cell system more directly relevant to the pathophysiology of human HHT, we generated blood outgrowth ECs (BOECs) from circulating endothelial progenitors in peripheral blood of an HHT patient genetically confirmed to carry the ALK1 T372fsX truncation (Fig. 6C). Strikingly, in these cells, tacrolimus robustly increased pSmad1/5/8 and ID1 levels, showing that tacrolimus promoted Smad1/5/8 signaling in HHT patient ECs (Fig. 6D).

**Figure 6.**
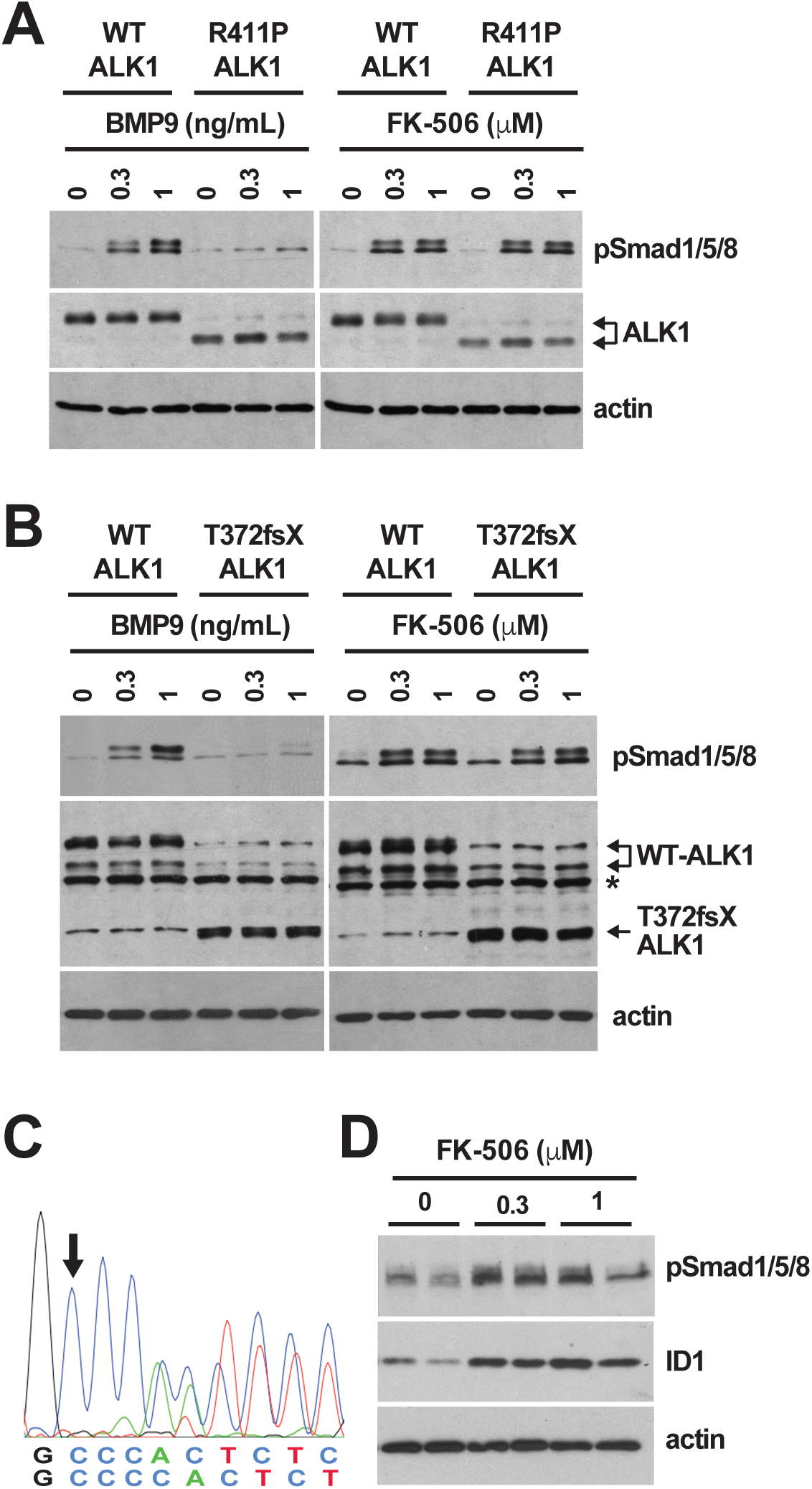
Tacrolimus activates Smad1/5/8 in cells expressing ALK1 HHT mutants. (**A and B**) C2C12 cells transfected with human WT ALK1 or ALK1 HHT mutants R411P (A) and T372fsX (B) were serum-starved for 3h (0.1% FBS medium) and then treated for 3h with the indicated concentrations of BMP9 or tacrolimus (FK-506). Asterisk denotes a nonspecific band. (**C**) Partial genomic DNA sequences of the HHT patient *ACVRL1* gene. Arrow indicates the c.1112dup insertion (T372fsX mutation) in the mutated allele. (**D**) HHT BOECs were treated with the indicated concentrations of tacrolimus for 24h in depleted medium (0.1% FBS medium). Cell extracts in (A, B, and D) were analyzed by WB using antibodies directed against the indicated proteins.

## Discussion

Whole-transcriptome analyses in primary ECs revealed that tacrolimus is a potent ALK1 signaling mimetic at the transcriptional level. Specifically, we report that tacrolimus treatment almost diametrically opposed the transcriptional deregulation caused by ALK1 inhibition by significantly changing in the reverse direction, the expression of 187 genes among the 251 found to be deregulated in the ALK1 loss-of-function transcriptional response (Fig. 1). Notably, ALK1 loss-of-function transcriptional response was associated with expression elevations of several key pro-angiogenic regulators, such as *DLL4*, *ANGPT2*, *KDR*, *CXCR4, PGF,* and *SMOC1*. Previous studies have demonstrated that BMP9/10-ALK1 signaling is required for the maintenance of vascular quiescence (43-45). Disruption of this quiescence mechanism is believed to facilitate the reactivation of angiogenesis and cause the appearance of abnormal vessels in HHT. In adult inducible *Alk1* and *Eng* knockout mice, for instance, wound-induced AVMs were reported to originate from angiogenesis-dependent vascular elongation (46, 47). In addition, extensive and concordant evidence from our laboratory (40) and others (19, 43, 48-52) has demonstrated using different models that the expression of *ANGPT2*, *KDR*, and *CXCR4* is negatively controlled by ALK1 signaling. The control of *DLL4* expression by ALK1 signaling has also been reported in an *alk1* mutant zebrafish model (53) and in BMP9/10-treated pulmonary arterial ECs (17). In line with these studies, we found clear evidence that, at steady-state, ALK1 inhibition in HUVECs increased Dll4 expression at the transcript and protein levels. We further show that Smad1/5/8 signaling activation with tacrolimus or BMP9 had the opposite effect by repressing Dll4 expression (Figs. 2 and 3). Together, these results further highlight the central role of ALK1 in the transcriptional regulation of angiogenesis and thus, fully support the concept that ALK1 loss-of-function and HHT pathogenesis are associated with deregulated activation of angiogenesis.

Because VEGF signaling is critical for Dll4 expression during angiogenesis, we asked whether tacrolimus impacts VEGF signaling activation. Our data in primary ECs showed that tacrolimus is a potent inhibitor of VEGF-mediated activation of Akt and p38 (Fig. 4). These results are in line with previous work showing that tacrolimus could interfere with VEGF-mediated EC tube formation *in vitro* (54). Because BMP9 treatment failed to recapitulate the repressing effect of tacrolimus on VEGF-mediated Akt/p38 activation, we concluded that the drug controlled VEGF signaling independent from its effect on ALK1 (Fig. 4C). We therefore postulate that tacrolimus lowered Dll4 expression *via* a dual control of ALK1 and VEGF signaling, two pathways critically, but independently, involved in Dll4 expression regulation. Although these data strengthen the concept that ALK1 cross talks with Notch signaling during angiogenesis, it is worth mentioning that studies performed in zebrafish have demonstrated that ALK1 can also regulate vascular quiescence and AVM development *via* a mechanism independent of Notch (53, 55). It will therefore be important going forward to determine the exact contributions of the different Dll4/Notch signaling pathways in AVM development in ALK1 loss-of-function models.

A recent study has demonstrated that BMP9/ALK1 blockade was associated with PI3K/Akt overactivation and that PI3K inhibition prevented the HHT vascular pathology in mouse models (56). In this context, our data suggest that tacrolimus treatment might also be beneficial by mitigating the aberrant Akt activation observed in HHT models. It is interesting to note that the authors of this study (56) also showed that, although ALK1 inhibition enhanced VEGF-mediated angiogenesis, its effect on Akt overactivation was, at least in part, not dependent on VEGF signaling. It appears therefore that ALK1 might control Akt signaling in ECs *via* several independent pathways. Future studies will be needed to assess the specific contributions of these different Akt pathways in the aberrant angiogenic process of HHT pathology and to determine whether tacrolimus affects their signaling responses differently.

An important question relates to the mechanism by which tacrolimus activates Smad1/5/8 signaling. FK-506-binding protein-12 (FKBP12) is known to interact with TGF-β type I family receptors within their receptor glycine-serine-rich phosphorylation domain to mute receptor activation in the absence of ligand (57, 58). More recently, compelling data showed that tacrolimus can activate BMPR2 and Smad1/5/8 signaling in pulmonary EC by blocking the inhibitory binding of FKBP12 to ALK1, but also to ALK2 and ALK3 (35), two other BMP type I receptors that are widely expressed. In this study, we confirmed that tacrolimus intervened upstream at the level of the receptor by activating Smad1/5/8 and by promoting a decrease of surface ALK1 in primary ECs, demonstrating that upon tacrolimus treatment, ALK1 responded to receptor activation through internalization (36).

Our data further demonstrated that tacrolimus was capable of potently activating Smad1/5/8 in cells overexpressing R411P- and T372fsX-ALK1, two well-described ALK1 HHT mutations that rendered the cells unresponsive to BMP9 (Fig. 6). These results are important because it was proposed that HHT pathology might arise from complete ALK1 loss-of-function (3). A drug with therapeutic potential is thus expected to activate Smad1/5/8 signaling in the presence of non-functional ALK1 receptors. In line with these results using transfected cells, we found that tacrolimus also potently activated Smad1/5/8 in BOECs isolated from an HHT patient carrying the ALK1 T372fsX truncation. These results not only show that tacrolimus activates Smad1/5/8 signaling in the presence of defective ALK1 receptor and in HHT patient ECs, but also are consistent with the proposed model that tacrolimus can control Smad1/5/8 signaling in ECs by bypassing ALK1 loss-of-function through the parallel activation of ALK2 and ALK3, for instance (35).

Treatment of newborn mice with BMP9/10 blocking antibodies leads to the development of a specific pathology in the developing retinal vasculature, which includes abnormal hypervascularization (17, 18), but also AVMs when ALK1 signaling is inhibited at a vascular developmental stage that allows a normal blood flow (40, 43-45). Recently, we reported that antibody delivery and BMP9/10 immunoneutralization in mouse pups could be achieved *via* the transmammary route by injecting lactating females. We further showed that this model is more practical, robust, and particularly suitable and reliable for the analysis of large cohorts (40). Tacrolimus administration in this model elicited a robust reduction in retinal hypervascularization and a small but significant decrease in AVM numbers and diameter (Fig. 5). Strikingly, tacrolimus also potently increased *Id1* levels in retinal ECs, while reducing the elevation of Dll4 levels observed in the BMP9/10-immunoblocked retinas. Immunohistochemistry analyses suggested that the Dll4 elevation triggered by BMP9/10 immunoblocking was restricted to the ECs located at the developing vascular front. Knowing that Dll4 is a tip cell marker (26), it is tempting to speculate that tacrolimus, by activating Smad1/5/8 signaling in this model, has anti-angiogenic properties that limits a pro-tip cell phenotype required for the hypervascularization observed in HHT models. Further investigation will be required to test this hypothesis and delineate the exact effect of tacrolimus on EC specification deregulations in HHT.

Finally, our data concur with previous studies that provide indirect evidence supporting the potential beneficial effect of tacrolimus in HHT patients. Indeed, a Canadian case study reported the regression of angiodysplasia and reduction of mucosal hemorrhage in a probable HHT patient who underwent liver transplantation following high-output cardiac failure and hepatic AVM development (59). The authors proposed that the anti-rejection regimen, which included tacrolimus, contributed to the observed beneficial effects in this HHT patient. In addition, the potential clinical benefit of tacrolimus in patients with end-stage PAH has recently been reported (60).

In conclusion, we show in this study that tacrolimus is as a potent activator of Smad1/5/8 signaling in cell and mouse models of ALK1 loss-of-function, including in HHT-patient derived primary ECs. Consequently, tacrolimus treatment corrected the signaling and gene expression defects caused by endothelial ALK1 inhibition and significantly improved HHT vascular pathology in mice. We propose that tacrolimus has therapeutic potential in HHT.

## Methods

### Cell cultures and transfections

HUVECs were isolated from anonymous umbilical veins and subcultured in 5% fetal bovine serum (FBS)-containing EC growth medium (Sciencell), as described before (40, 61). C2C12 cells were obtained from ATCC. C2C12 cells expressing pNeo-(BRE)2-Luc reporter plasmid (C2C12BRA) were kindly provided by Dr. Daniel Rifkin (34). C2C12 cells were maintained in Dulbecco’s Modified Eagle’s Medium (DMEM) supplemented with 10% FBS, penicillin and streptomycin. Cells were transiently transfected with pcDNA3 plasmids encoding C-terminally HA-tagged WT ALK1 or ALK1 HHT mutants T372fsX (also annotated G371fsX391) and R411P [kindly provided by Dr. Sabine Bailly (14)], using Lipofectamine 2000 reagent (Invitrogen) and as per the manufacturer’s instructions.

### Human blood outgrowth endothelial cells (BOECs) isolation and characterization

BOECs were isolated from blood draws obtained from an HHT2 patient carrying the ALK1 T372fsX truncation [*ACVRL1* c.1112dup mutation, first reported in Ref. (42)]. Cells were isolated as described before (62). Flow cytometry was performed to characterize the generated BOECs (data not shown), using the surface markers: VEGFR2 and CD31 (endothelial), and CD45 (hematopoietic, negative control), following a procedure described in (62). BOEC genomic DNA was isolated using DNeasy blood and tissue kit (Qiagen), as per the manufacturer’s recommendations. *ACVRL1* exon 8 partial sequence was amplified by PCR using forward primer ACTCACAGGGCAGCGATTAC and reverse primer CCAAAGGCCCAGATGTCAGT. The generated 142 pb PCR product was then sequenced using reverse primer AAAGGCCCAGATGTCAGTCC.

### RNA extraction and sequencing

HUVECs were rinsed with PBS and processed for RNA extraction using the RNeasy mini kit (Qiagen), according to the manufacturer’s instructions. Total RNA quality was verified using Thermo Scientific NanoDrop and Agilent Bioanalyzer. RNA was processed for RNA-Seq at the Genomics Resources Core Facility, Weill Cornell Medical College, New York, NY. Briefly, cDNA conversion and library preparation were performed using the TrueSeq v2 Illumina library preparation kit, following manufacturers’ recommended protocols. Samples were multiplexed 6 per lane and sequenced on an Illumina HiSeq 4000 instrument.

### RNA-Seq data analysis

Reads were uploaded to the GobyWeb system (63) and aligned to the 1000 genome human reference sequence (64) with the STAR aligner (65). Ensembl annotations for transcripts were automatically obtained from Biomart and Ensembl using GobyWeb. Annotations were used to determine read counts using the Goby alignment-to-annotation-counts mode (66), integrated in GobyWeb in the differential expression analysis with EdgeR plugin. Counts were downloaded from GobyWeb as a tab delimited file and analyzed with MetaR (67). Statistical analyses were conducted with Limma Voom (68), as integrated in MetaR 1.7.2, using the rocker-metar docker image version 1.6.0.1. *P*-values were adjusted for multiple testing using the False Discovery Rate (FDR) method (69). Heat maps were constructed with MetaR, using the pheatmap R package. Gene annotations were determined with Ensembl/Biomart, using the biomart micro-language in MetaR (67). MetaR analysis scripts are presented in Supplementary Material (Figs. S2-S4). UpSet plot were generated with MetaR and the UpSet R package (70).

### Drug screening and luciferase assays

BMP9 EC_50_ was determined in C2C12BRA cells following treatments with different concentrations of recombinant BMP9 for 24h in depleted medium containing 0.1% FBS. FDA-approved drug libraries (NIH clinical collections, NCCs) were screened at 3 μM on C2C12BRA cells maintained for 24h in depleted medium containing 0.1% FBS and 0.5 ng/mL BMP9 (BMP9 EC_50_). Cells were solubilized in Luciferase Cell Culture Lysis 5X Reagent (E1531, Promega) and processed for luciferase measurements using Luciferase Assay System (E1501, Promega), following manufacturer’s instructions and using Victor^3^ 1420 Multilabel counter luminometer (PerkinElmer).

### Mice

Timed-pregnant C57BL/6J mice (3-4-month-old, The Jackson Laboratory) were used in this study.

### Transmammary-delivered immunoblocking of BMP9 and BMP10, tacrolimus treatments, and retinal vasculature analyses in mice

Antibody injections and retinal whole-mount histochemistry were performed in mice as previously described (40). Briefly, lactating dams were injected i.p. once on P3 with mouse monoclonal isotype control antibodies (15 mg/kg, IgG2b, MAB004; 15 mg/kg, IgG2a, MAB003; R&D Systems) or mouse monoclonal anti-BMP9 and anti-BMP10 antibodies (15 mg/kg, IgG2b, MAB3209; 15 mg/kg, IgG2a, MAB2926; R&D Systems, respectively). On P3, P4, and P5, pups were injected i.p. with tacrolimus (FK-506, 0.5 mg/kg) or vehicle (1% DMSO saline). Pups were euthanized on P6 by CO_2_ asphyxiation and were enucleated. Eyes were fixed in 4% paraformaldehyde for 20 min on ice and retinas were isolated and analyzed by histochemistry, as before (40), see also *Supplementary Methods*. Images for the analysis of the vascular network density were acquired using a laser confocal microscopy Olympus FV300. Quantifications were performed using ImageJ. Using a 20x lens, images (2-5 fields per retina) were acquired at the vascular plexus (between an artery and a vein). Quantification was done by using the measure particles tool, working with 8-bit images, adjusting the threshold, and measuring the area occupied by the vasculature in a region of interest of 200 x 200 μm^2^.

### Retinal EC isolation and RT-qPCR

ECs were isolated from neonatal retinal tissue using anti-CD31 microbeads (Miltenyi Biotec GmbH), as per the manufacturer’s instructions. One-step RT-qPCR was performed on pellets of 40-60,000 cells per sample using Cells-to-Ct^TM^ 1-Step TaqMan^®^ Kit following the manufacturer’s protocol (Ambion, Thermo Fisher Scientific Inc.). PCR was performed using TaqMan assays on ABI7900HT (Applied Biosystems, Life Technologies). *Id1* expression levels were normalized to the reference gene *Polr2a*. Relative changes in gene expression were determined by the ΔΔCt method (71) and using control values normalized to 1.0.

### Statistics

Figure legends indicate the test used for each experiment to determine whether the differences between the experimental and control groups were statistically significant. A *P* value < 0.05 was considered to be statistically significant. The analyses were performed using GraphPad Prism 7 (GraphPad Software Inc.).

### Study approvals

Study subject (HHT2 patient carrying the ALK1 T372fsX mutation) provided voluntary and written informed consent using a form approved by the Feinstein Institute for Medical Research Institutional Review Board (IRB). Study subject BOECs were isolated and cultured using a protocol approved by the Institute’s IRB. All animal procedures were performed in accordance with protocols approved by the Feinstein Institute for Medical Research Institutional Animal Care and Use Committee, and conformed to the NIH Guide for the Care and Use of Laboratory Animals and ARRIVE guidelines (72), see *Supplemental Methods*.

## Author contributions

S.R., P.C., H.Z., J.P., P.K.C., and P.M. performed experiments and analyzed data. F.C. supervised the RNA-Seq and performed the bioinformatic analyses. E.C. assembled the IRB application and performed the subject blood draws. P.M. conceived the project and elaborated the experimental strategy with S.R., C.N.M., L.B., and F.C.; P.M., S.R., F.C., L.B., and C.N.M. wrote the manuscript.

## Data and materials availability

RNA-Seq reads have been deposited to the Sequence Read Archive (accession number pending).

## Acknowledgments

We thank Dr. Daniel Rifkin (NYU School of Medicine, New York, NY) for providing us with C2C12BRA cells and Dr. Sabine Bailly (CEA Grenoble, France) for the ALK1 plasmids. We are grateful to Drs. LaQueta Hudson and Kevin J. Tracey (The Feinstein Institute for Medical Research) for assistance with the NIH clinical collections.

## SUPPLEMENTARY MATERIAL

### SUPPLEMENTARY METHODS

#### Materials and antibodies

Recombinant human ALK1-Fc (Cat. # 770-MA), BMP9 (3209-BP), and VEGF_165_ (293-VE) were obtained R&D Systems. Tacrolimus was from Cayman Chemical (10007965). For flow cytometry, antibody directed against ALK1 was from R&D Systems (AF370). For Western blots, antibodies directed against pSmad1/5/8 (13820), Smad5 (9517), p-Erk1/2 (9101), Erk1/2 (9102), p-p38 (4511), p38 (9212), p-T308-Akt (2965), p-S473-Akt (4060), Akt (9272) were obtained from Cell Signaling Technology; anti-ID1 antibody from BioCheck (BCH-1/195-14); anti-actin antibody from BD Transduction Laboratories (612656); anti-Dll4 antibody from R&D Systems (AF1389); anti-HA antibody from Roche (3F10, used to detect transfected HA-tagged ALK1 in Fig. 6A); anti-ALK1 antibody from Santa Cruz Biotechnology (sc-101555, used to detect transfected ALK1 in Fig. 6B). For immunohistochemistry (IHC), anti-Dll4 antibody was obtained from R&D Systems (AF1389) and donkey anti-goat Alexa Fluor 633 secondary antibody from Molecular Probes (A21082). PCR primers were obtained from IDT and sequencing was performed at Genewiz.

#### Western blot (WB) analyses

Cells were processed as before (1) with the following modifications. Cells were solubilized in RIPA buffer (20-188, EMD Millipore) supplemented with 1× Complete protease inhibitor mixture (11697498001, Roche). 5-20 μg of proteins (depending on the primary antibody used) were separated by SDS-PAGE and transferred onto nitrocellulose membranes. Membranes were then probed with primary and secondary antibodies. A standard ECL detection procedure was then used.

#### Flow cytometry

HUVECs were stained with anti-ALK1 extracellular domain antibody AF370 (20 μg/mL final concentration) for 15 min at room temperature (RT). After 2 washes in PBS 0.5% (w/v) BSA, cells were incubated with a donkey anti-goat Alexa Fluor 488 (1:2000 dilution) for 15 min at RT. Cells were washed twice in PBS 0.5% BSA and fluorescence was measured using a BD Fortessa cytometer. Subsequent analyzes were performed using FlowJo software v10.0.8.

#### Mice

To conform to ARRIVE guidelines (2) and control for potential bias, animals were randomized by arbitrary selecting half of each litter to go into the control or tacrolimus treatment group. Researchers who assessed the treatment outcomes (S.R. and P.M.), i.e., by retinal vascularization analyses, were blinded to mouse allocation and treatment, which were performed by H.Z. No allocation concealment, however, was used. Based on sample size calculations, mixed groups of male and female mice (litters of 7-9 pups) were treated on P3 (average weight 2 grams) with anti-BMP9/10 antibodies or vehicle (PBS) using the transmammary route and as described in details in Ref. (3). The same mouse groups were also treated daily with tacrolimus (0.5 mg/kg) from P3 to P5 and sacrificed on P6 for analyses. Mice were maintained in regular housing conditions and were allowed free access to water and maintenance diet.

#### Retinal whole-mount histochemistry

After fixation using 4% paraformaldehyde, retinas were dissected, cut four times to flatten them into petal flower shapes, and fixed with methanol for 20 min on ice. After removing methanol, retinas were washed in PBS for 5 min on a shaker at RT, and incubated in blocking solution (0.3% Triton, 0.2% BSA in PBS) for 1h on a shaker at RT. Retinas were then incubated in isolectin GS-IB4 Alexa Fluor 488 (I21411, Molecular Probes) diluted 1:100 in blocking solution on a shaker overnight at 4°C. Retinas were then washed four times in 0.3% Triton in PBS for 10 min on a shaker, followed by two washes in PBS for 5 min on a shaker before mounting with Vecta Shield (H-1000, Vector Laboratories). For Dll4 IHC, 5% donkey serum was added to the blocking solution, and an additional incubation step with donkey anti-goat Alexa Fluor 633 secondary antibody (1:2000, Molecular Probes) was performed.

---

**Figure S1.**
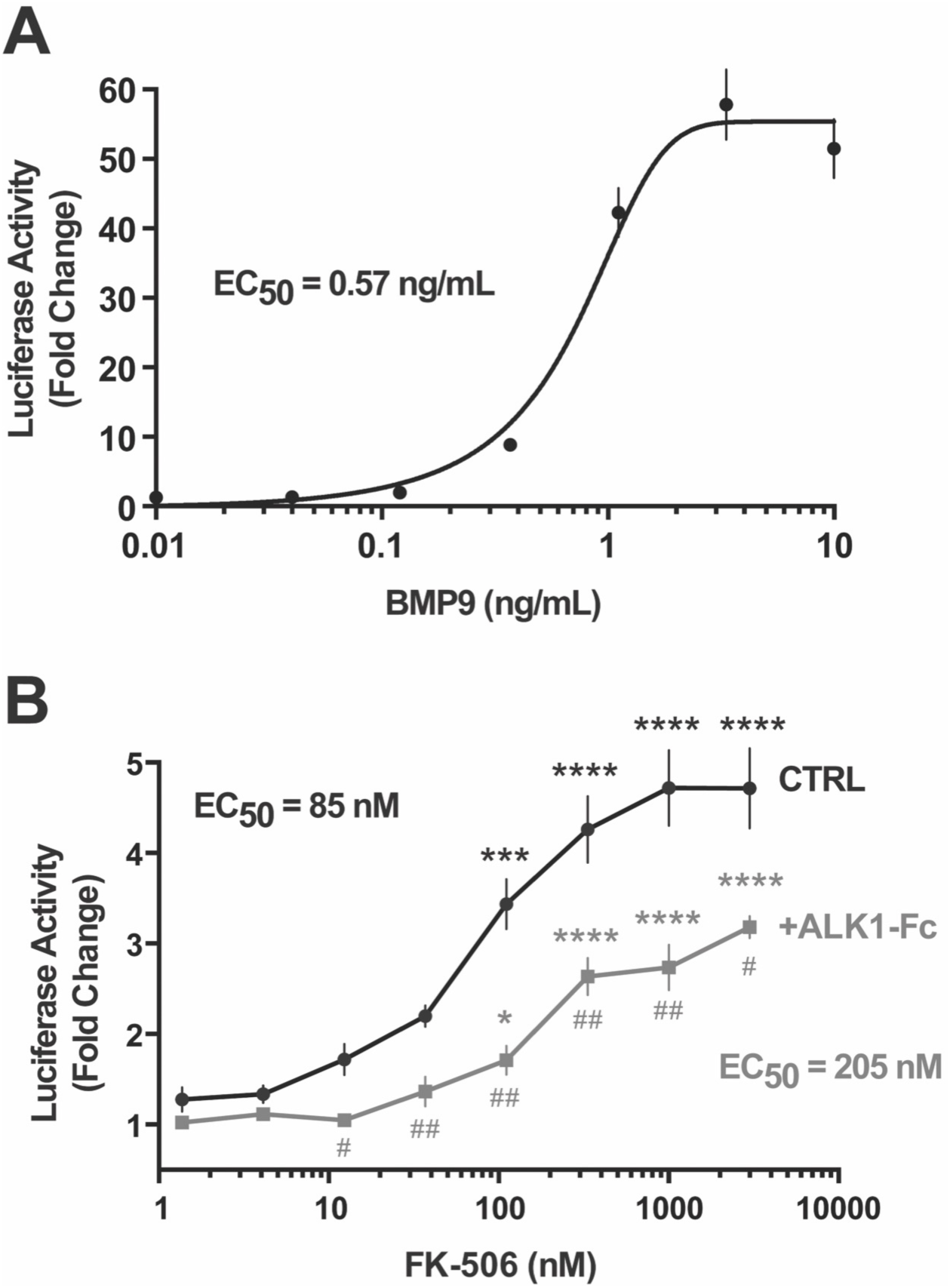
BMP9 and tacrolimus dose-responses in C2C12BRA cells. (**A**) Luciferase activity in C2C12BRA cells treated with different concentrations of BMP9 for 24h in depleted medium (0.1% FBS). Data represent mean ± s.e.m. (n = 5). (**B**) Luciferase activity in C2C12BRA cells treated or not (CTRL) for 24h in complete medium (conditioned for 2 days) with ALK1-Fc (1 μg/mL) and the indicated concentrations of tacrolimus (FK-506). Data in (A) and (B) represent mean ± s.e.m. (n = 4); \**P* < 0.05, \*\*\**P* < 0.001, \*\*\*\**P* < 0.0001; one-way ANOVA, Dunnett’s multiple comparisons test; ^#^*P* < 0.05, ^##^*P* < 0.01 relative to non-ALK1-Fc-treated controls at the corresponding FK-506 concentration, Student’s *t*-test.

**Figure S2.**
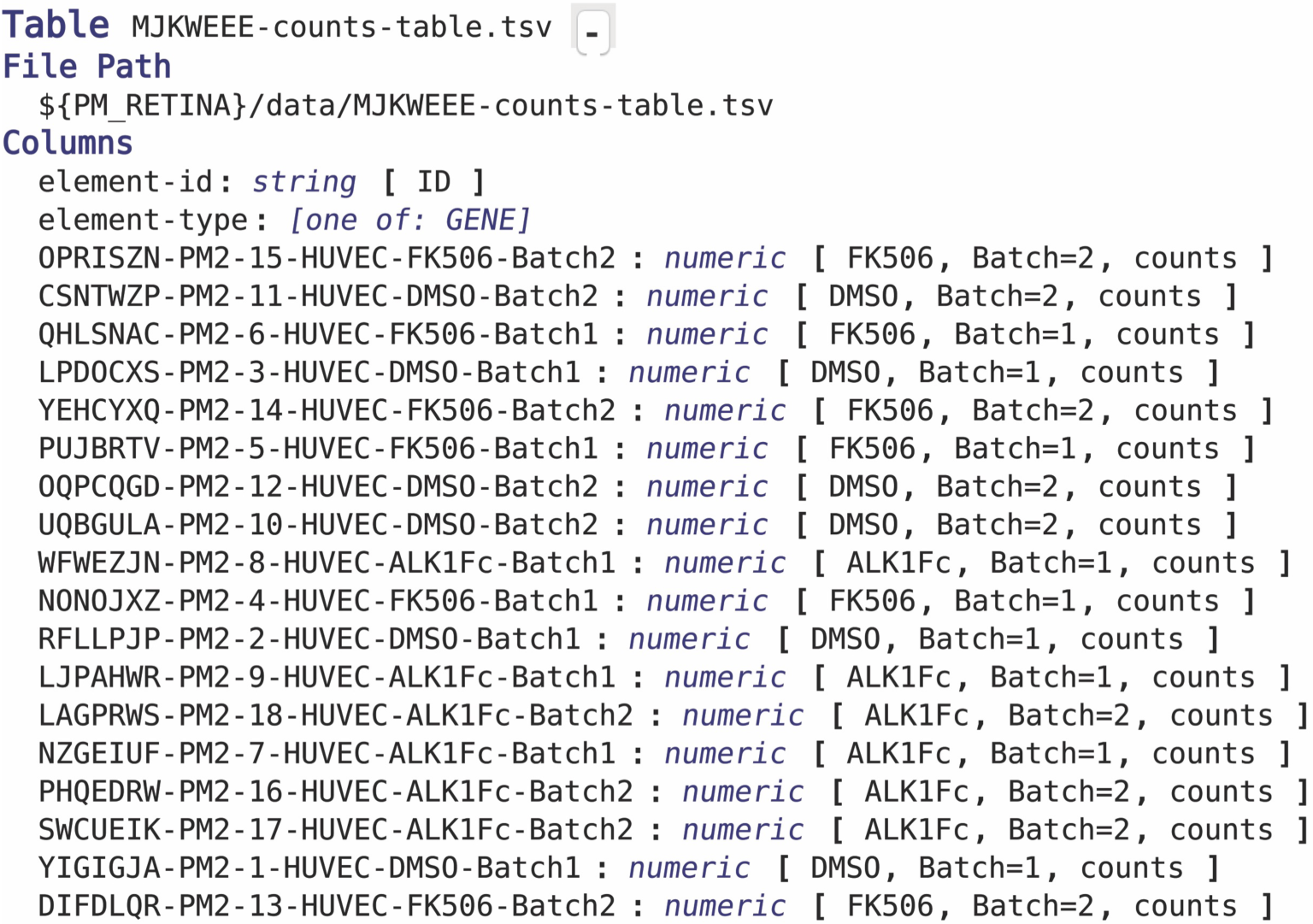
Table annotation for MetaR Analysis.

**Figure S3.**
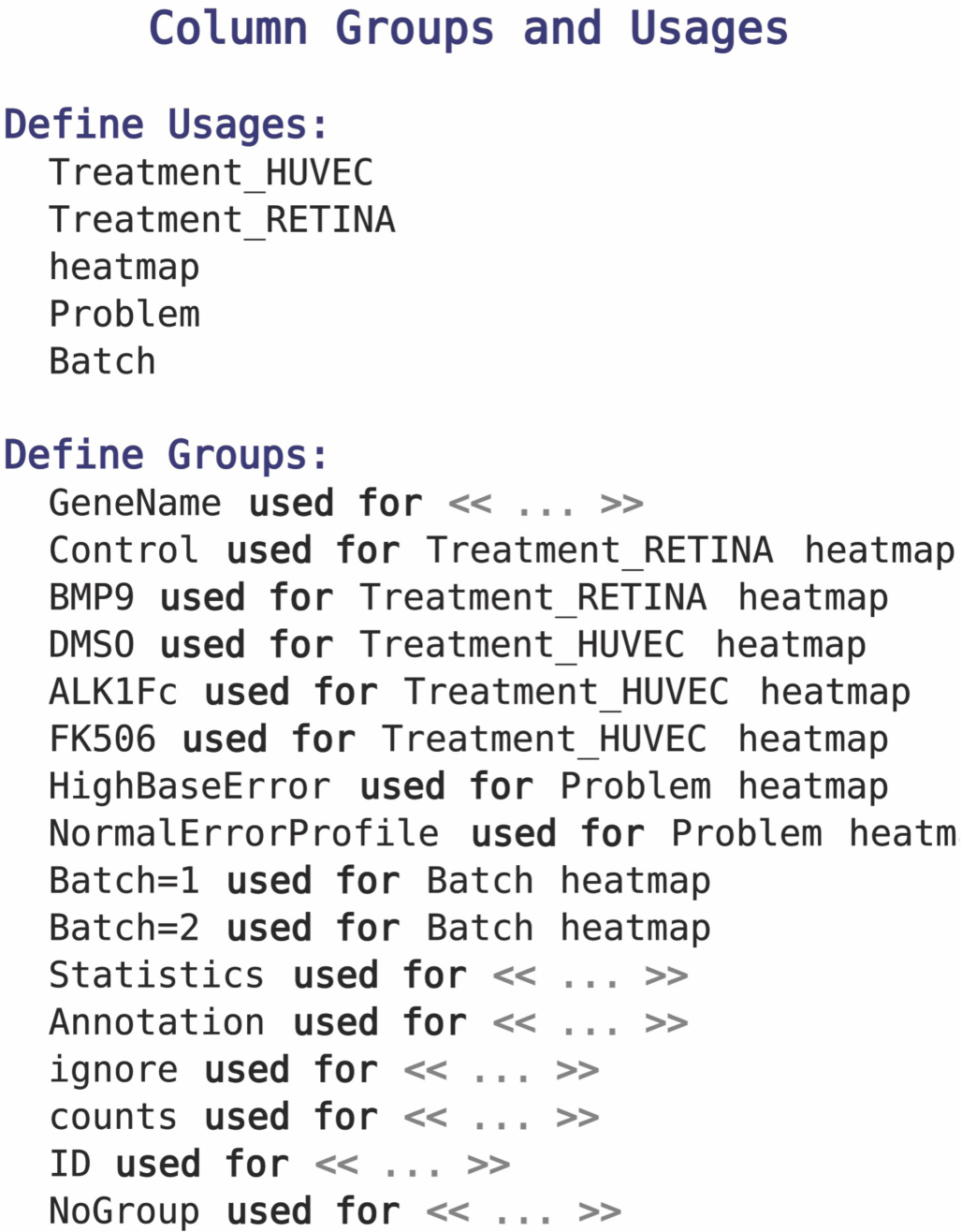
Group definitions for MetaR Analysis.

**Figure S4.**
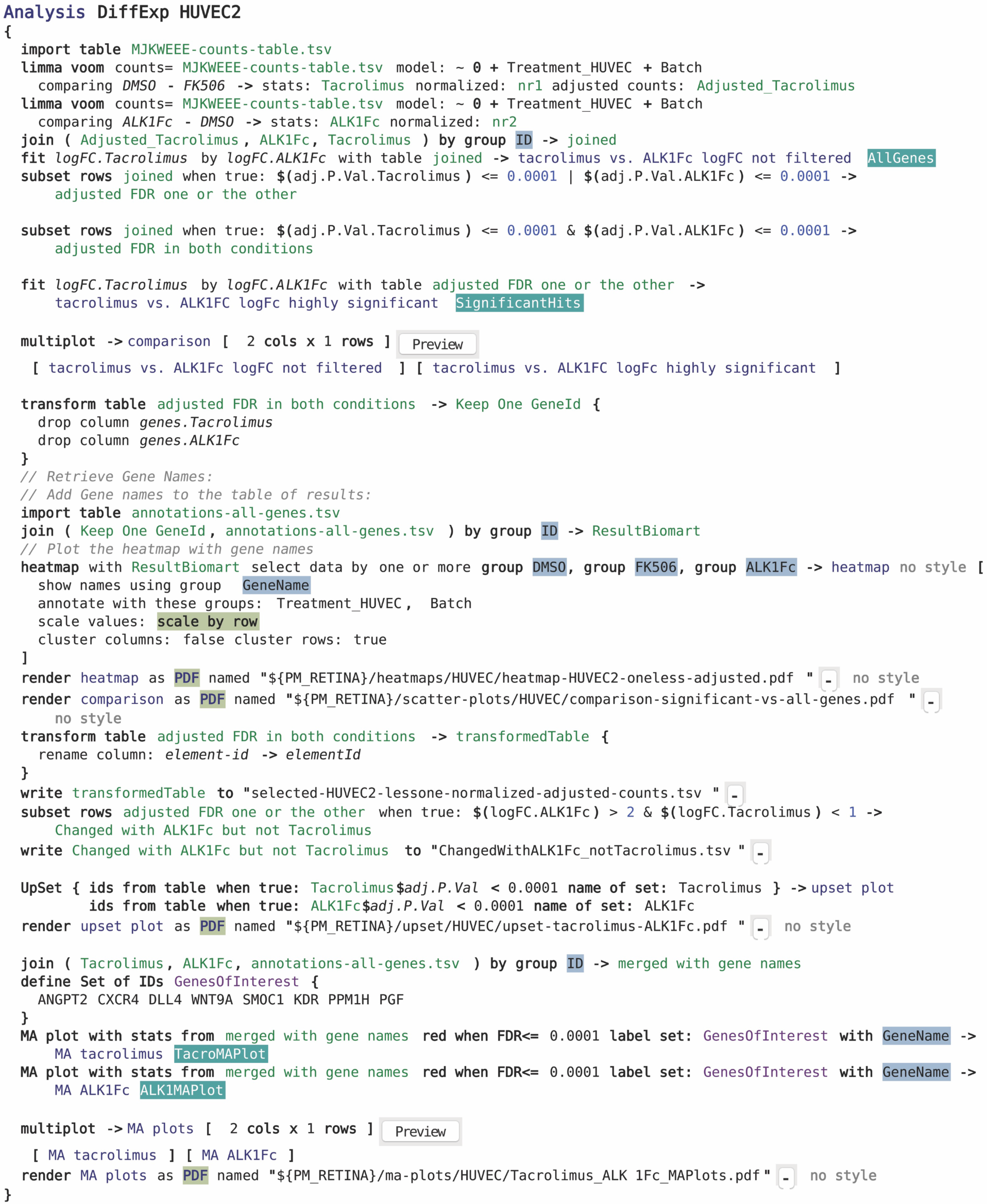
MetaR analysis script for RNA-Seq analysis.

**Table S1. RNA-Seq analyses in HUVECs.** Gene expression changes were analyzed in HUVECs treated or not (DMSO) for 24h in complete medium (conditioned for 2 days) with ALK1-Fc (1 μg/mL) or tacrolimus (FK-506, 0.3 μM). Table presents the annotated results (selected-HUVEC2-lessone-normalized-adjusted-counts.tsv, imported into Excel, produced by the analysis script shown in Fig S4) and describes differentially expressed genes with a FDR less or equal to 0.0001 (n = 5-6).

